# The ratio versus difference optimization and its implications for optimality theory

**DOI:** 10.1101/2020.12.15.422822

**Authors:** Sonali Shinde, Ankur Patwardhan, Milind Watve

**Affiliations:** Department of Biodiversity, Abasaheb Garware College, Pune, India; Independent Researcher, Pune, India

**Keywords:** Optimality theory, ratio optimum, difference optimum, Concorde fallacy, parental investment, r and K selection, pollination, behavioral economics

## Abstract

Among the classical models of optimization, some models maximize the ratio of returns per investment, others maximize the difference between returns and investment. However, the question under what conditions use of the ratio model is appropriate and under what conditions a difference model should be used remained unaddressed until recently. The question is important because the strategies indicated by ratio optimum can be substantially different than the ones suggested by difference optimum. We make a general case here for the set of conditions for appropriate use of ratio versus difference optimum. When the investable amount is perceived as limiting but not the investment opportunities a ratio optimum is appropriate and when the investment opportunities are perceived to be limiting but not the investable amount, difference optimum is appropriate. Taking examples of Concorde fallacy, parental investment, r and K selection, nectar production, pollinator behavior, protein synthesis and stability, viral burst size and human economic behavior we show that the ratio-difference distinction in optimization models resolves many long standing debates and conundrums in evolution, behavior and economics.

## Introduction

Optimality theory is an important element of behavioral ecology and evolutionary psychology which assumes that a strategy that optimizes the cost-benefits gets selected. Optimization models improve our understanding about adaptation and innate behavioral tendencies (Parker & Maynard-Smith 1990). A number of examples of behavioral optimization abound literature in behavioral ecology (Stephens & Kerbs 1987; Ramírez-Bautista *et al*., 2000; Thakar *et al*., 2003; Ha 2010; Doniol-Valcroze *et al*., 2011). Although there has been serious criticism on some aspects of optimality theory (Pierce & Ollason 1987), the criticism is mainly about empirical testing and practical applications of optimality theory. It is really tough to have a null model and a sound experimental design for claiming that the observed behavior of a species is optimal in a given context. Further there is lack of convergence in our understanding of evolutionary or adaptive optimization on the one hand and the psychological, neurobiological or proximate mechanisms at work on the other (Stevens 2008). Fewer studies attempt convergence of proximate and ultimate causes in optimization (Budaev *et al*., 2019; Baig *et al*., 2019). Nevertheless, at a conceptual level, optimization theory continues to be useful to address evolutionary, behavioral and economics questions (Rahnev & Denison 2018). Not all well studied examples of optimization come from species believed to have high levels of intelligence. Complex optimization decisions have been modeled and studied in insects (Abe & Kamimura 2012; Charnov & Skinner 1988; Mills & Zaviezo 2000; Tenhumberg *et al*. 2006) and bacteria (Watve *et al*., 2006; Lele *et al*., 2011; Baig *et al*., 2014). Further, optimization principles originated in biology have been used to develop useful and efficient algorithms (Chen *et al*., 2011). Complex organisms exhibit much behavioral plasticity and have evolved the ability to judge the cost benefits in a given context and optimize their behavior contextually. Behavioral optimization models have been used to explain human behavior in nutritional (Nettle *et al*., 2017), ecological (Watve *et al*., 2016) and social context (Purshouse & McAlister 2013). There is considerable debate over the application of optimality to humans (Driscoll 2009; Rahnev & Denison 2018) particularly in experimental tests of perceptual decision making. However, even in these tests although individuals deviate substantially from the optimum, the average performance of a group is close to the theoretical optimum (Rahnev & Denison 2018). Despite limitations, constraints and contextuality, optimization is a useful fundamental concept in behavioral evolution.

However, there are certain fundamental questions in the optimality theory that remain unaddressed. In a landmark review of optimization models, Parker and Maynard-Smith (1990) discussed many examples of optimization models in some of which the difference between investment and returns is maximized and in others the ratio of the two is maximized (Figure 1). However, why it is appropriate to use the ratio in some examples and difference in others is not explained. For single unit investment optimization, although both ratio models and difference models are used in literature, the question when a ratio model is appropriate and when a difference model is appropriate was not formally addressed until recently (Watve *et al*., 2016., Watve & Ojas 2020). We argue here that the outcomes of ratio versus difference optimization can be substantially different; clarity in the conditions under which ratio or difference optimization is appropriate resolves many long standing questions and conundrums; and this distinction gives new clarity about some of the old debates.

**Figure 1:**
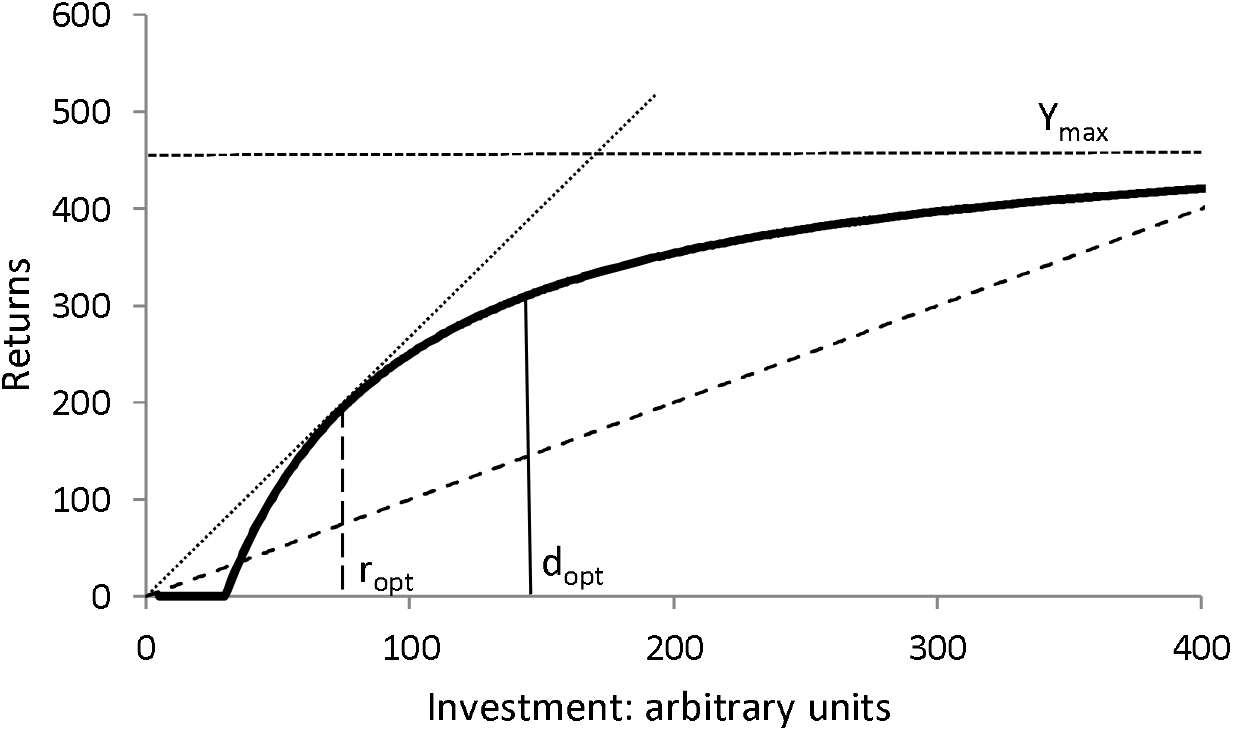
A conceptual diagram explaining ratio optimization and difference optimization: When the output follows the law of diminishing returns and there is an overhead cost, the inputs where the benefit-cost ratio is maximized is where the tangent drawn from the origin touches the curve. The benefit-cost difference is maximized where the vertical difference between the curve and the break even line is maximum, or where the slope of the curves is unity. For any profitable relationship the ratio optimum is always on the left of the difference optimum. How much effort one invests in a behavioral act depends upon whether ratio is being maximized or difference is being maximized.

## The model

We begin with a baseline assumption of a saturation curve, following the law of diminishing returns which is the baseline assumption of most optimization models in behavioral ecology (Parker and Maynard-Smith 1990), economics (Brue 1993) and anthropology (Foley 1985), despite certain contextual differences across disciplines. Conceptually the differentiation between ratio and difference optimization can be used with other shapes of curves as well but we will use the saturation curve to develop a baseline model.

The intuitive logic behind optimization with a saturation curve having a non-zero overhead cost is quite simple. Graphically the ratio optimum lies where the tangent drawn to the curve from the origin touches the curve (Figure 1). The deal is profitable when the slope of this tangent > 1. The difference optimum lies where the distance between the curve and the break even line is maximum, or where the slope of the curve =1. Since the slope of the curve is always decreasing, the difference optimum will always lie to the right of the ratio optimum provided the curve allows a profitable deal. The ratio optimum can be exactly equal to the difference optimum only when the tangent has a slope =1, i.e. the best possible deal is only a break-even.

A difference optimization model maximizes the benefit per investment opportunity and therefore, when investment opportunities are limiting, this is the model of choice. On the other hand, the ratio model maximizes the benefit per unit investment and therefore when the investable amount is limiting, a ratio optimum should be used (Figure 1). In reality, an individual can have multiple investable units and the net benefit of an individual is the sum of benefits from all invested units. There are two possible scenarios in which the total or lifetime benefit can be maximized.

### A. Optimizing when complete information about the investable amount and investment opportunities is available at a time

If information about the investable amount, the number of investment units and the benefit curve is available, an optimum investment per unit which will maximize the total returns can be worked out as follows.

If *T* is the total investable amount one has and *N*, the maximum number of units available for investment, one should choose the number of units *n* for investment and *C* the amount invested in each unit to get a return *Yc* per unit such that, *Y_c_.n+T-n.C* is maximized. The actual number of units invested in and the amount invested in each unit hold a constrained relationship such that n <= N/C. Assuming a saturation curve within each unit described by the equation

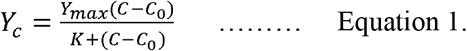

we choose *C* that maximizes *Y_c_.n+T-n.C* with the above constraint on *n*. Using numerical simulations we get typical curves (figure 2) in which the optimum investment per unit transits between a lower limit and an upper limit. It can be seen that at the lower limit, the ratio of *Yc/C* is maximum and at the upper limit the difference *Yc-C* is maximum. Thus the two limits of the optimum investment per unit correspond to the ratio optimum and difference optimum. When *T* is limiting but not *N*, the ratio optimum gives maximum net returns summed over all investment units. On the other hand when *N* is limiting but not *T*, difference optimum gives maximum net returns. There is only a brief transient range when the true optimum lies between the two limiting optima. Calculating an optimum in the transition zone needs prior information on all parameters.

**Figure 2:**
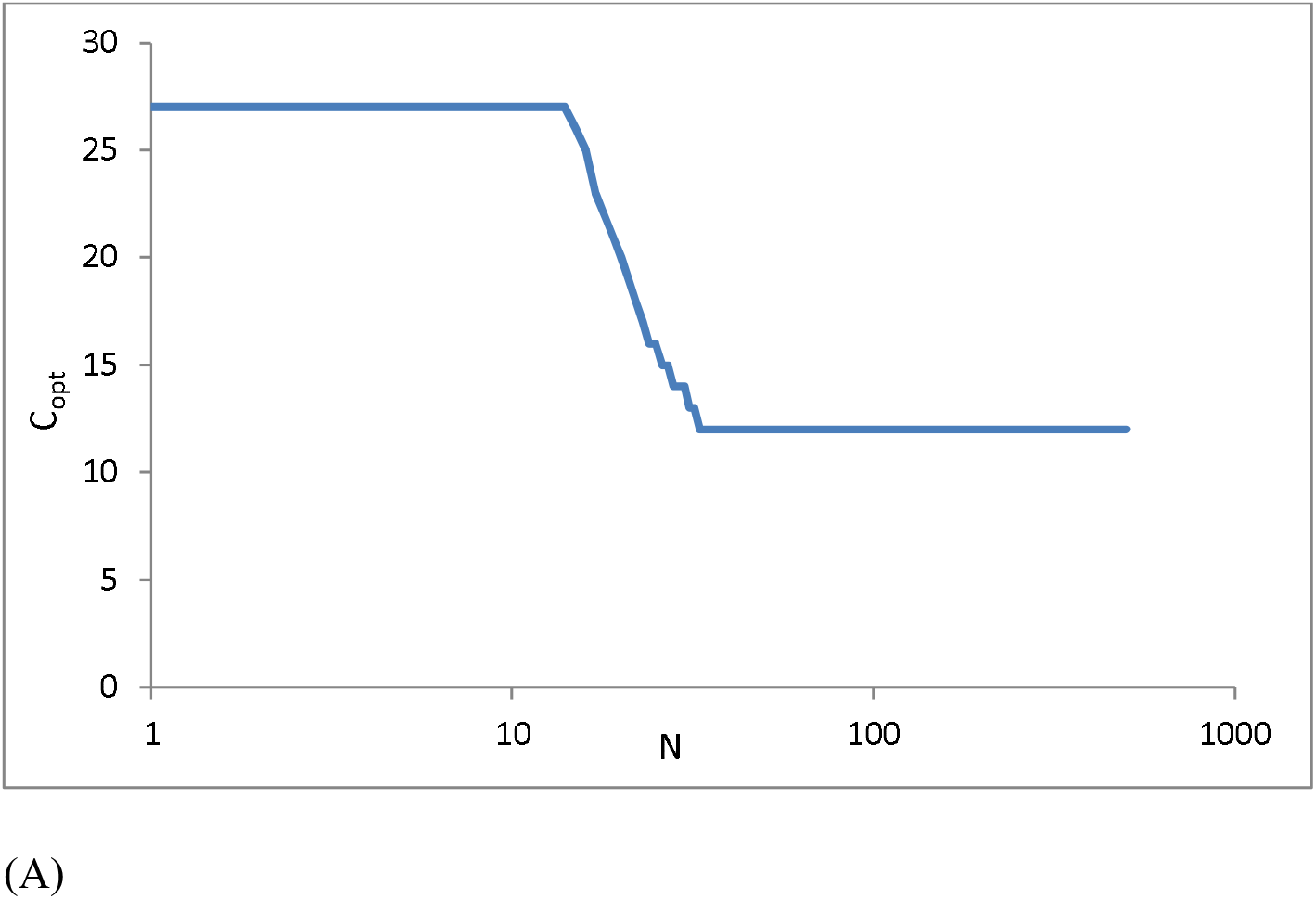
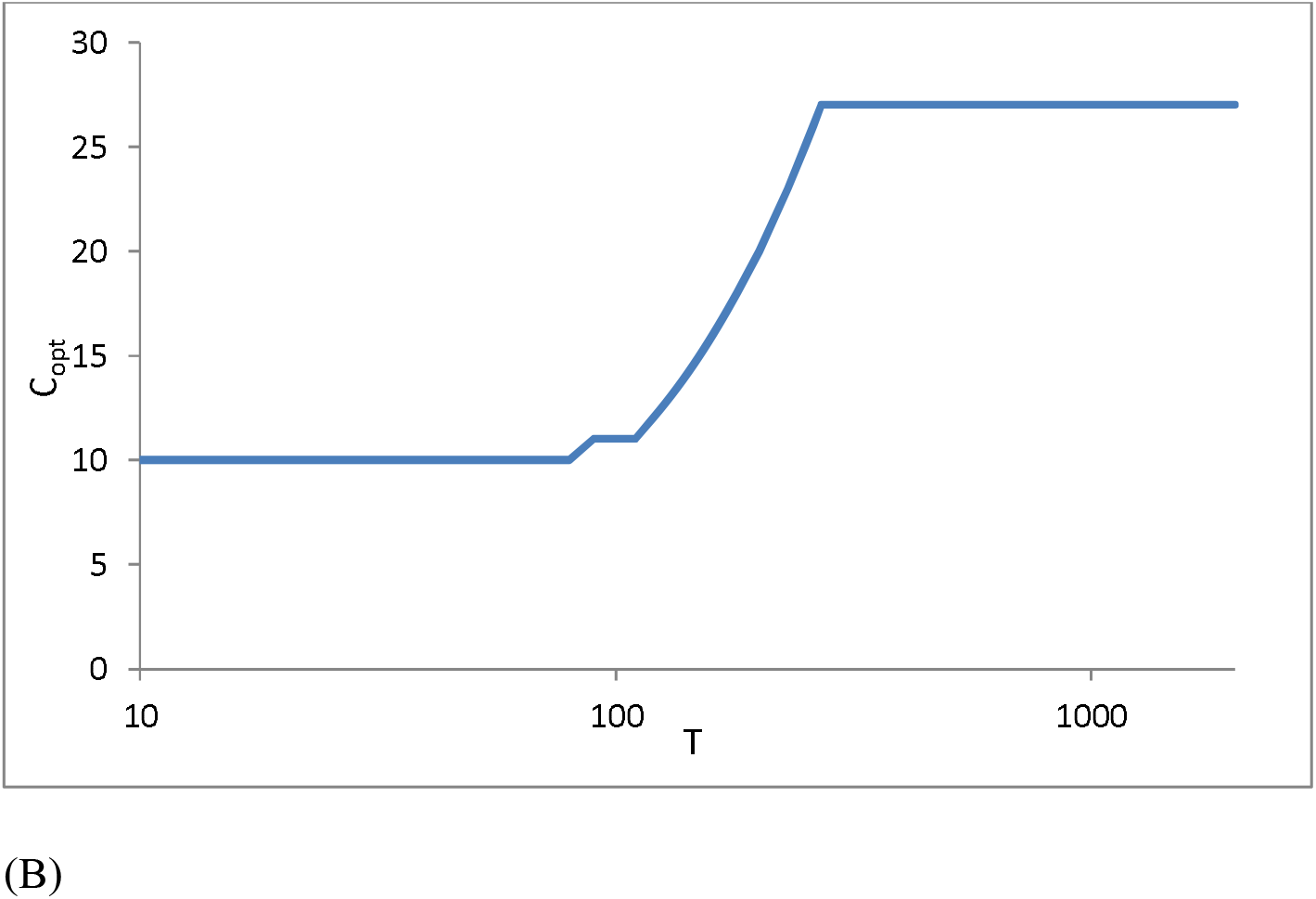
A. The optimum investment per unit with a fixed T and variable N. The optimum has two plateaus with a brief transition zone. When N is limiting there is a higher plateau and when N becomes so large that T becomes limiting, there is a lower plateau. B. Similarly with constant N and variable T you see a lower plateau when T is limiting and a higher plateau where N is limiting. There is a brief transition zone when neither T nor N is clearly limiting.

Going by equation 1, for ratio optimum 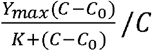 is maximized and for difference optimum, 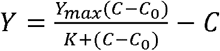 is maximized. By taking derivative *dY/dC* and finding *C* when the derivative becomes zero (Watve *et al*., 2016), the optimum investment to maximize the ratio can be shown to be,

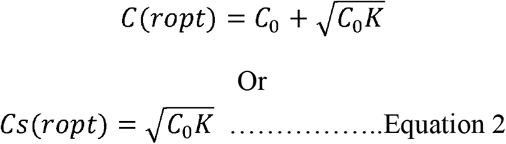

**Where** *C_s_*=(*C – C*_0_).

And the difference optimum to be,

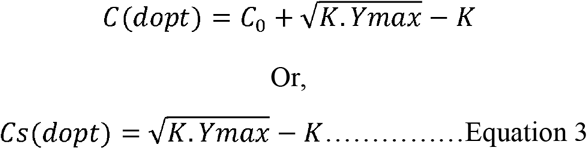

When *C*_(*dopt*)_. *N <T*, the investable amount is not limiting but the number of investment units is. In this case difference optimum should be used and when *C*_(*ropt*)_.*N > T*, the investable amount becomes limiting and not the investment opportunities. In this case ratio optimum should be used. However since *C_(dopt)_* > *C_(ropt)_* there will be a condition when *C_(ropt)_.N < T <C_(dopt)_.N*

This condition is the transition zone between the ratio and difference optimum as in figure 2. When *T* is in this range and for a given *Cs* the net profit is

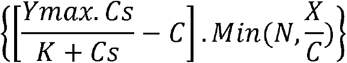

*Cs* at which the net profit is maximized will be the optimized *Cs*. A typical curve during transition from ratio to difference optimum is as in Figure 3.

**Figure 3:**
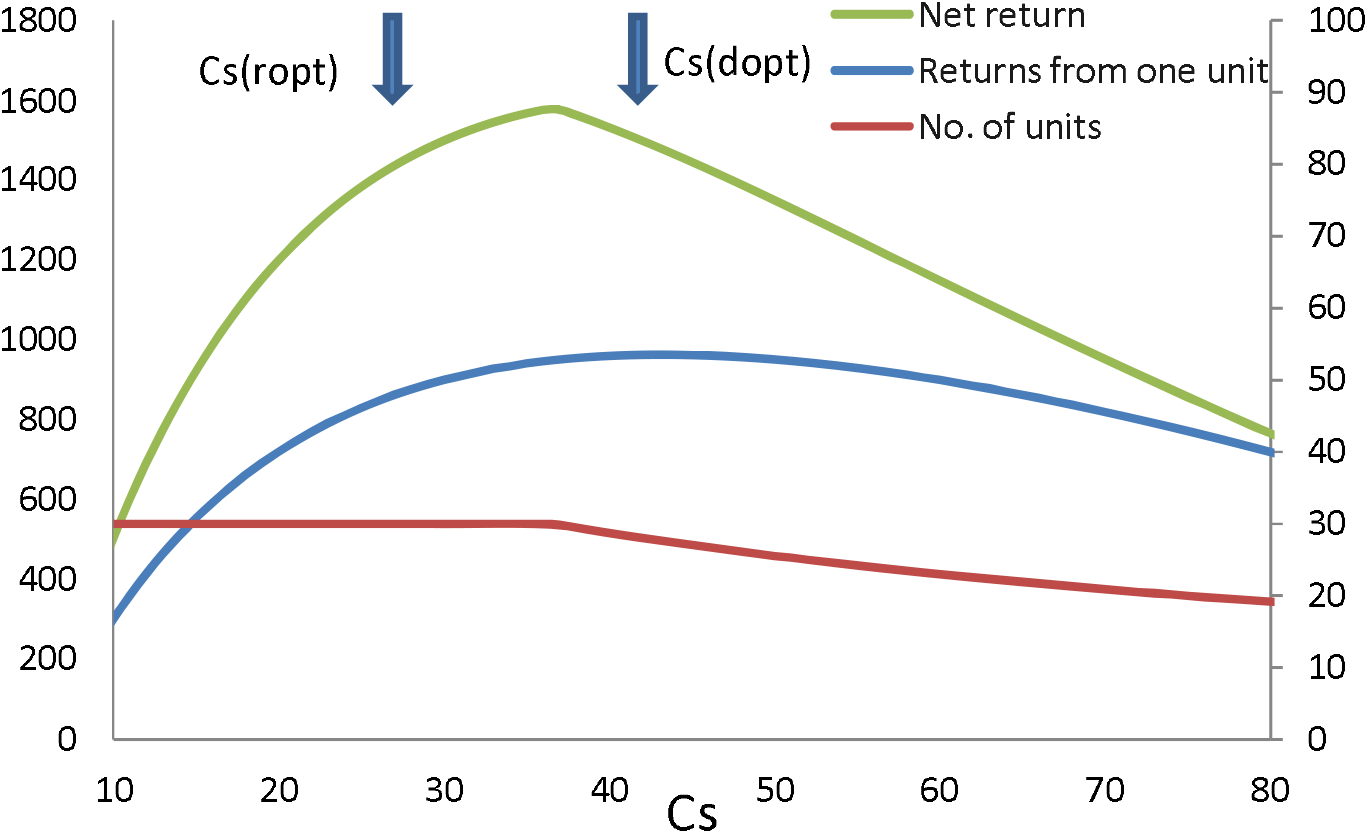
When C_(ropt)_.N < T < C_(dopt)_.N, i.e. neither the investable amount nor the opportunities are clearly limiting, the optimum Cs lies between the ratio optimum and difference optimum. Other parameters used here are C_0_=40, Y_max_=200, K=20, N=30 and T =2300.

It can be easily seen that for finding the optimum in the transition zone knowing the parameters of the curve is not sufficient. One needs complete information about *T* and *N* simultaneously.

### B. Optimizing with incomplete information

With real life constraints, complete information about the lifetime investable amount or the number of investment opportunities available in lifetime is hardly ever available. These variables may also change with time so extrapolative calculations are also unreliable. Therefore, most decisions are likely to be based on optimizing the investment in one unit at a time. With incomplete information a simple rule of thumb would be to go by the ratio optimum or the difference optimum depending upon whether the perceived limiting factor is *T* or *N*. This is the most likely scenario in practice and therefore we expect most innate optimization methods as well as cognitive decisions to be based on either of the two limiting optima.

From equation 2 and 3, it can be seen that optimum *C_s_* for the ratio model increases with increasing overhead cost, but is independent of *Y_max_*. The optimum *C_s_* for the difference model is independent of the overheads, but varies with the square root of *Y_max_*. These two distinctions can be used as differential testable predictions of the two models. Ratio optimizers should increase their investment when the overheads increase. Difference optimizers’ optimum running cost should be independent of the overheads. A change in the maximum possible profit, i.e. *Y_max_* on the other hand, will change a difference optimizer’s investment in proportion to the square root of *Y_max_*, whereas that of a ratio optimizer should remain unaltered.

We will see below how a strategic decision to use the ratio optimization or difference optimization influences the decision making in human and non-human examples.

## Applications and implications

The clarity about using ratio or difference optimization has many implications to behavioral ecology, evolution, economics and policy making. We intend to explore only a few such examples in which the ratio difference concept is likely to bring better clarity and offer novel solutions to long standing riddles. There is extensive background literature discussing all these issues and we neither intend to systematically review or critique it, but only suggest which new dimension or clarity can be added to these concepts by the ratio-difference optimization model.

1. Concorde fallacy: Concorde fallacy or sunk-cost is a long standing conundrum. Even when it was clear that the Concorde airplanes would not bring any profit, the British and French airlines continued using them on the grounds that they had spent a large amount of money on recruiting them, which should not go waste. To discontinue using them would have avoided further net losses and therefore to continue using them is termed economically irrational behavior. There are examples of Concord fallacy or sunk cost fallacy in animal as well as human behavior the reasons for the origin and survival of this behavior is a conundrum with a wide variety of attempts to explain (Arkes & Ayton 1999; Baliga & Ely 2011; Roth *et al*., 2015; Doody 2020). We propose here that the apparent paradox can be understood based on the ratio versus difference model. An airline company can potentially recruit any number of planes. So the limiting factor is not the number of planes but the amount that the company decides to invest. If the deal was perceived to be profitable, the ratio model would have been the appropriate strategy. Note that the optimum investment in a ratio model is independent of *Y_max_* (Figure 4). So when the actual revenue curve turned out to be much lower than the one projected, for ratio thinkers the intended duration of use would not change. The difference optimum, on the other hand would reduce substantially if the curve fails to rise as expected. Therefore a difference optimizer would advise termination of use as soon as the slope of the curve becomes less than unity. The ratio optimizer, on the other hand would continue till the originally projected use. So the continued use of Concorde in spite of absolute loss is not entirely irrational. It’s a different consideration. If a ratio model is indeed appropriate for the context, then it is not a fallacy. However, if a ratio model is used when it should have been a difference model, it is indeed a fallacy. The animal and human examples of sunk cost behavior need to be re-examined for the appropriateness of the ratio model for the context. It is likely that we are interpreting the decisions assuming difference models and the lack of clarity between the contextual appropriateness of the two models makes it appear as a fallacy.
2. Male and female parental investment: Sexual selection, sexual dimorphism, adult sex ratio, parental care and the difference in male female investment strategies are intertwined concepts and the causal relations between them have multiple interpretations (Trivers 1972; Clutton-Brock 1991; Kokko & Jennions 2008; Fromhage & Jennions 2016; Jennions 2017; Ratikainen 2018). By the fundamental biology of sex, females have greater investment in offspring than males mainly in terms of egg formation and maternal care wherever applicable. The males often have large investment in secondary sexual character, attractive display and the risk associated. Inclusive of this, males’ average total reproductive investment divided over all his offspring may not be less than the female average. Therefore, unlike classical assumption, the difference in male and female strategies is not explained by the total investment per offspring. The ratio-difference optimization brings in much clarity in the complex dynamics of parental investment. Since the number of offspring a male can potentially have is not limiting, males are expected to be ratio optimizers. Females cannot increase the number of offspring by mating with multiple males and therefore they are limited by the investment opportunities and therefore are difference optimizers. Therefore, even if the total reproductive investment per offspring of males and females might turn out to be the same, their investment strategies need not be the same. The male’s investment in secondary sexual character and display is an overhead cost in this model, but unlike our baseline model, males have a dual overhead cost. There is a preparative overhead necessary before access to females which is in the form of investment in size, strength and/or display infrastructure. This component of investment does not increase with the number of females courted or mated. We call this type 1 overhead. The type 2 overhead is the additional cost to be paid per every female courted and consists of courtship and other behaviors targeted specifically and separately to individual females and/or individual offspring. Only type 2 overheads can be considered as *C_0_* of the model going by the model assumptions. Therefore although the male’s total investment may not always be lower than a female’s, owing to the two stage overheads and use of ratio model, male’s effective investment per offspring is more likely to be smaller than that of the female (Figure 5). If offspring survival is enhanced by paternal investment, the female should try to increase the overhead type 2 of the male by seeking more intensive courtship since in ratio optimization, the optimum *C_s_* increases with increase in *C_0_*. A testable prediction of the model is that in species without the need for paternal care, type 1 overhead should dominate male displays but in species where some paternal inputs are needed, female choice and sexual selection will favor type 2 overheads, i.e. active and individual oriented courtship over general display characters. Exercising a choice for type 2 overheads may result in an overhead cost for the female as well, but since in difference optimization, the optimum *C_s_* is independent of the overhead, maternal investment per offspring will not be affected. There are two types of factors affecting parental investment according to the ratio-difference optimization. A parent’s optimum investment can change if the investment-returns curve changes. What type of change in the curve brings about increase or decrease in the investment would depend upon whether a ratio or difference optimization is being used. For example greater predatory pressure can reduce offspring survival and thereby *Y_max_* of the curve. We have seen that the ratio optimum is independent of *Y_max_* whereas the difference optimum is dependent. We expect therefore that female investment per offspring would depend upon the presence or absence of predator but male investment may not. The male investment, on the other hand will increase if the type 2 overheads increase, but female investment will not. These principles can be further used to make species specific testable predictions of the ratio-difference concept in the parental investment theory. Our generalization that males are expected to be ratio optimizers and females difference optimizers may change in specific contexts. If the number of surviving offspring is severely opportunity limited, even for a male, difference optimization may become the model of choice and males would invest more in parental care (Benowitz *et al*., 2013). Therefore a parent’s investment strategy may change either due to a change in the model itself or due to a change in the characteristics of the curve. The two interact with each other and the nature of their interaction can also be modeled on the platform of the ratio-difference principle.
3. <u>The quality-quantity trade-off in offspring</u>: The theory of r and K selection perceived the trade-off between the number of offspring and the investment per offspring (McArthur and Wilson 1967; Reznick *et al*., 2002). Some aspects of the r and K selection theory came under criticism later and a more elaborate life history optimization theory was proposed (Pianka 1970; Gadgil & Bossert 1970; Michod 1979; Kozlowski 1980; Charlesworth 1980; Stearn 1976, 1977, 1992; Derek 1993; Vitzthumb, 2008). The life history optimization also talks about optimum investment per offspring. It is recognized that selection for quality versus quantity of offspring is under a diversity of selective forces (Wilbur 1974; Cassill 2019). The concept of ratio versus difference optimization can complement these theories and bring in greater clarity. Under certain contexts the number of offspring is limited by parental investment and in certain other contexts by environmental opportunities. When the environmental opportunities are in excess of what parental investment permits, the ratio optimum should decide the investment per offspring and when the environmental opportunities are limiting the difference optimum should be chosen. It can be seen that under a set of assumptions, the ratio optimum is mathematically equivalent to r selection and difference optimum to *K* selection. If *T* is the lifetime investment in reproduction, *C/T* is the number of offspring. Taking *T=1*, the birth rate,

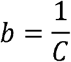 If *Y* is interpreted as survival probability of the offspring with *Y_max_* <=1, *1-Y* is the death probability and *b(1-y)* the death rate. Therefore

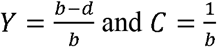 Maximization of ratio, 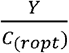 is equivalent to maximization of 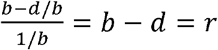 Maximization of difference *Y – C*_(*dopt*)_ is equivalent to maximization of

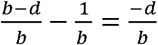 Maximizing −*d/b* is maximizing *b/d* which is equivalent to *K*. Kozolowski (1980), Maynard Smith (1989), Charnov (1986) and Stearns (1992) agree that *r* selection maximizes *b-d* and *K* selection maximizes *b/d*. Therefore the ratio optimization is mathematically equivalent to r selection and difference optimization to *K* selection. The classical *r* and *K* selection concept talks about two alternative directions of selection but does not specify how far selection will drive along these directions, whereas the ratio versus difference optimum depicts two distinct optima. Where exactly *b – d* is optimized and where *b/d* is given by the cost-benefit models. This is likely to resolve at least some of the ambiguities associated with the r and K selection theory (Kozolowski 1980).
4. Optimizing the proportion of nectarless flowers: It is known that in any species of flowering plants a proportion of flowers contain nectar while others are empty (Simpson *et al*., 1981; Bell 1986; Thakar *et al*., 2003; Anand *et al*., 2007). There are multiple models of optimization of the proportion of the empty or “cheater” flowers (Bell 1986; Southwick 1987; Pyke 1991; Charnov *et al*., 2006; Bailey *et al*., 2007; Morris *et al*., 2010; Thakar *et al*., 2003; Belasare *et al*., 2009). The models so far have not explicitly considered alternative scenarios of whether the availability of pollinators is limiting or the number of flowers that can be made. If pollinators are limiting, the plants should follow a difference optimization model. In areas with pollinator abundance, they should follow ratio optimization. Thakar et al (2003) and Belasare et al (2009) used a difference model without explicitly defining the limiting factor. Incorporating the ratio-difference optimization in these models is likely to give better insights as well as make differential testable predictions that can drive novel experimental designs. The counter-part of the nectar optimization by the plant is the optimization of foraging strategy by the pollinator (Waddington 1979; Smithson and MacNair 1997; Smithson A and Gigord 2001; Sidarus *et al*., 2019). This would depend upon whether flower availability and spatial distribution is limiting or available time is. Accordingly the optimization strategy of the pollinator would change.
5. Optimizing viral burst size: For a virus infecting unicellular or multicellular hosts the time for which a given cell is used for replication needs to be optimized. There can be a variety of rate limiting steps in this process (Cummings *et al*., 2012; Yin *et al*., 2018). Initially the virion population within a cell replicates exponentially as long as the virion number itself decides the rate of growth as seen in bacteriophage (Berngruber 2013). At a later stage, however, some component of the cell’s replication machinery can become limiting. Therefore a virus has a choice to burst out early before the machinery becomes rate limiting, to maximize the reproduction per unit time. However, by utilizing a cell for a longer time, a virus can maximize replication per cell, but replication per unit time would be lower. Depending upon whether the constraint is time or the availability of cells, the length of the cycle and burst size would be optimized. For example, the influenza virus can invade a body only until the acquired immune response develops sufficiently to eliminate it (Chen *et al*., 2018). Since time is the constraint, the virus needs to evolve a faster cycle with small burst size rather than compromising on the rate of growth for a longer life cycle with a larger burst size. The virus can afford to make sub optimum use of the cell capacity to support viral replication, because being a respiratory virus, finding another host is easier. In contrast, in the case of HIV, immune response is impaired on the one hand but on the other the mode of transmission being specialized and restrictive (Iwasaki, 2012), finding another host is more difficult. Therefore influenza virus should be a ratio optimizer investing small time per cell and HIV should be a difference optimizer investing more time and utilizing the replication potential per cell fully. Compatible with the expectation is the estimated burst size of the slow growing HIV to be about 50000 (Chen *et al*., 2007) and that of the faster growing influenza virus to be 500-1000 (Mahmoudabadi *et al*., 2017).
6. Protein stabilization: For a cell, proteins need to be synthesized and then folded and maintained in the appropriate configuration (Cooper, 2000).The cell has an investment in protein synthesis and another set of investment in maintenance that includes chaperons (Bukau *et al*., 2006; Tang *et al*., 2008), and antioxidant mechanisms (Murray, 2018). In the worst case, the option of protein degradation and recycling of amino acids also exists. The investment in stabilizing a protein can be optimized either by the ratio model or the difference model. Baig et al (2014), showed that bacterial cells accumulate misfolded proteins when the external supply of nutrients is not limiting. On the other hand when nutrient supply is limiting, there is greater investment in protein maintenance and recycling. This strategic shift can be explained by ratio versus difference optimum. When protein synthesis is not limiting, the investment in protein maintenance is optimized by ratio. Therefore there are greater chances of proteins misfolding and aggregation. In contrast when nutrient resources limit protein synthesis, there is greater investment in protein maintenance and recycling. This is likely to be the strategic reason for some of the beneficial effects of caloric restriction in minimizing pathologies associated with protein aggregation (Baig *et al*., 2014; Matai *et al*., 2019; Yang & Zhang 2020)
7. Human behavior and policy decisions:

a. Mother’s investment in offspring: In a rural Ethiopian community, technological intervention to reduce the physical stress of mothers was expected to increase the health status of mothers along with improved child health. However, in reality, fecundity and birth rate increased leading to further worsening of child nutrition (Gibson & Mace 2006). A simplest and most appropriate explanation is offered by our model. In a community of ratio optimizer parents, if an intervention reduces mothers’ efforts in day to day work, it is equivalent to reducing the overheads in reproduction. Reducing overheads in a ratio model reduces the optimum investment per unit. Mothers in this community appear to have done the same. The researchers in this study suspect that birth control along with the intervention would have improved investment in child health. This means that the optimization model would have shifted from ratio to difference.
b. The economics of agriculture versus animal keeping: Watve & Ojas (2020) pointed out that in traditional sustenance agriculture practices, a farmer typically possesses only one farm. This is a context in which the investment opportunities are limiting. Therefore a farmer should use a difference optimum. In contrast, in traditional animal keeping, where animals are grazed in a common grazing land, the number of animals are unlikely to be limiting (Hardin 1968). Therefore, a ratio model is more appropriate. If animals are allowed to breed naturally, the overhead cost is also small. Therefore traditional animal keepers using common grazing grounds are expected to be ratio optimizers. This might be reflected in the differential response to hybrid seeds versus crossbred cattle in India. Both involve increased output but at a higher cost. The difference optimizing farmers accepted the high cost high returns practice. Animal keepers, being ratio optimizers, were keener to keep the denominator small and therefore gave a cold response to cross-breeding and artificial insemination programmes (Watve and Ojas 2020). The economics of animal keeping changes with private pastures/ranches. If the owner has sufficient investment capacity and the animals can only be grazed in his own land, the Pasture land becomes the limiting factor. Limited and exclusive pasture land puts an upper limit on the number of animals and makes it an opportunity limited case. So for private ranches a difference model is appropriate. In this model, animal keepers are keener to invest more in animals but expect greater returns. By this model we expect that the productivity per animal will increase with privatization of pasture land. In an open grazing system the total productivity will depend more on animal numbers than on productivity per animal. A testable prediction of the model is that selective breeding for high productivity animals is boosted by privatization of pasture lands.
c. Sustainable collection of seasonal natural resources: For people living in natural habitats and depending on multiple natural resources for livelihood, sustainable harvest is important for long term stability of livelihood resources. In biodiversity rich areas, multiple resources are available and the availability is often season dependent. We expect the success of collection to follow a saturation curve with input efforts. For a given resource, the crucial question is when to stop harvesting. If there are multiple options of livelihood, the decision would be ratio based. If there are few alternatives for livelihood, the decision would be difference based. A ratio based decision spares a greater amount of resources for regeneration. A difference based decision is likely to result in overharvesting. The tragedy of the commons (Hardin 1968) is more likely to happen if the habitat has fewer options or if the society has specialized communities monopolizing different resources. The latter is seen in many societies such as the traditional Indian endogamous communities with niche partitioning. For such communities overharvesting is more likely to be a potential hazard. However, since the community has little alternatives for livelihood, they need to assure sustainability and this is often achieved by making prudent harvesting norms for the community (Gadgil 1985; Gadgil & Malhotra 1985). Although the dynamics of natural resource harvesting has been a focus of investigation, the ratio-difference distinction in optimization has not been a part of the conceptual framework in this field. Studying the behavior of different communities dependent on natural resources in the light of ratio-difference strategies is likely to be both challenging and insightful.

**Figure 4:**
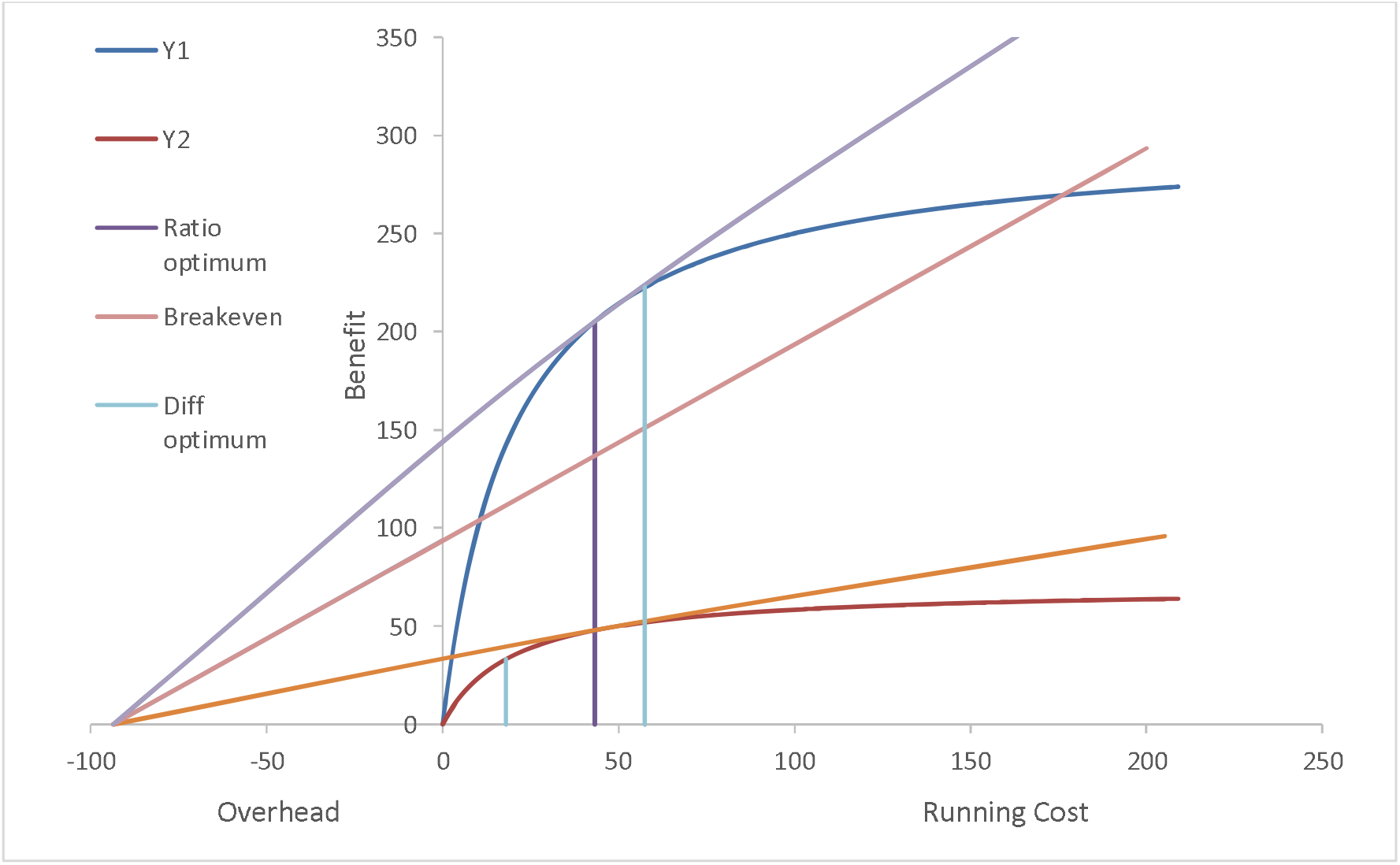
The Concorde fallacy viewed as ratio versus difference optimization model: The blue curve is the hypothetical projected revenue curve and the brown one the realized actual revenue collection. When the curve is flattened, the difference optimum shifts to the left considerably, but the ratio optimum does not change its position. Therefore a difference optimizer will stop wherever the slope of the curve <1, but the ratio optimizer will continue till the projected extent of use.

**Figure 5:**
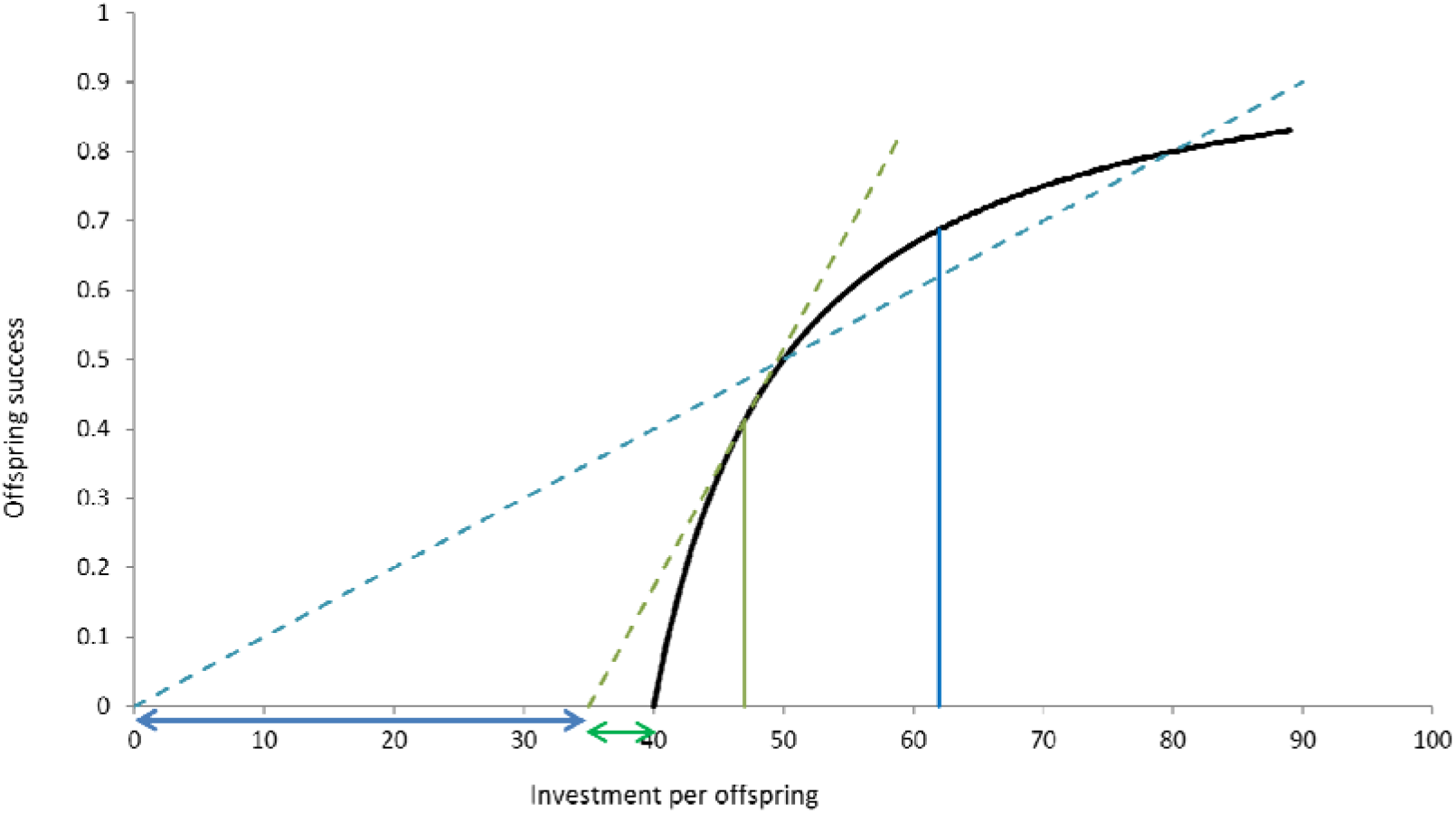
Difference between male and female investment in the offspring: We assume a case in which the net investment per offspring may be the same for males as well as females. We assume that the total overhead cost as well as the shape of the curve is the same. However, the nature of the overhead and the optimization models for males and females are different. The male overhead can be divided into the preparatory overhead (type 1-blue bar) and the courtship overhead (type 2-green bar). The former, once incurred, applies for all mating attempts and therefore does not figure in optimization of input per offspring. Secondly, males optimize the ratio. Females, on the other hand, have an overhead separately for every offspring and they optimize the difference. Therefore male investment in offspring, post fertilization is expected to be smaller than female investment, when other factors remain the same.

From the examples discussed, it can be seen that the ratio versus difference dichotomy can have implications for understanding the behavior of a variety of species or systems. The differentiation can address some long standing paradoxes and conundrums, develop new insights into problems as well as stimulate new questions and novel experimental designs.

